# CD73 maintains hepatocyte metabolic integrity and mouse liver homeostasis in a sex-dependent manner

**DOI:** 10.1101/2020.10.08.328930

**Authors:** Karel P. Alcedo, Morgan A. Rouse, Gloria S. Jung, Dong Fu, Marquet Minor, Helen H. Willcockson, Kevin G. Greene, Natasha T. Snider

**Affiliations:** Department of Cell Biology and Physiology; Department of Pathology and Laboratory Medicine, University of North Carolina at Chapel Hill

**Keywords:** adenosine, inflammation, AMP-activated protein kinase, ecto-5’-nucleotidase, hepatic zonation

## Abstract

**Background & Aims:** Metabolic imbalance and inflammation are common features of chronic liver diseases. Molecular factors controlling these mechanisms represent potential therapeutic targets. One promising target is CD73, the major enzyme that dephosphorylates extracellular adenosine monophosphate (AMP) to form the anti-inflammatory adenosine. In normal liver, CD73 is expressed on pericentral hepatocytes, which are important for long-term liver homeostasis. The aim of this study was to determine if CD73 has non-redundant hepatoprotective functions.

**Approach & Results:** We generated mice with a targeted deletion of the CD73-encoding gene (*Nt5e*) in hepatocytes (CD73-LKO). Deletion of hepatocyte *Nt5e* resulted in approximately 70% reduction in total liver CD73 protein (p<0.0001). Male and female CD73-LKO mice developed normally during the first 21 weeks, without significant liver phenotypes. Between 21-42 weeks, the CD73-LKO mice developed spontaneous onset liver disease with significant severity in male mice. Notably, middle-aged male CD73-LKO mice displayed hepatocyte swelling and ballooning (p<0.05), inflammation (p<0.01) and variable steatosis. Female CD73-LKO mice had lower serum albumin (p<0.05) and elevated inflammatory markers (p<0.01), but did not exhibit the spectrum of histopathologic changes characteristic of the male mice, potentially due to compensatory induction of adenosine receptors. Serum analysis and proteomic profiling of hepatocytes from male CD73-LKO mice revealed significant metabolic imbalance, with elevated blood urea nitrogen (p<0.0001) and impairments in major metabolic pathways, including oxidative phosphorylation and AMP-activated protein kinase (AMPK) signaling. There was significant hypo-phosphorylation in AMPK substrate in CD73-LKO livers (p<0.0001), while in isolated hepatocytes treated with AMP, soluble CD73 induced AMPK activation (p<0.001).

**Conclusions:** Hepatocyte CD73 supports long-term metabolic liver homeostasis through AMPK in a sex-dependent manner. These findings have implications for human liver diseases marked by CD73 dysregulation.

---

The highly integrated metabolic activities of hepatocytes control physiological homeostasis and are perturbed in chronic liver diseases. Non-alcoholic fatty liver disease (NAFLD) is the most prevalent chronic liver disease in children and adults, and a major risk factor for the development of cirrhosis and hepatocellular carcinoma (HCC) (1, 2). Lack of approved therapies renders NAFLD a major health problem, estimated to affect 25% of the global population, including 80 million people in the United States (3). In recognition that NAFLD is principally a metabolic disease, a new nomenclature was proposed: Metabolic-Associated Fatty Liver Disease (MAFLD) (4). From that standpoint, understanding how hepatocytes maintain long-term energy homeostasis will be critical for addressing the key mechanisms behind this complex and heterogeneous disorder.

It was shown over 50 years ago that ATP injections increase hepatic ATP content and can prevent the development of fatty liver in rodents (5). Moreover, patients with non-alcoholic steatohepatitis (NASH), the most severe form of NAFLD, have significantly impaired capacity for replenishing hepatic ATP stores after transient depletion (6). Numerous biochemical reactions control ATP metabolism and utilization by hepatocytes, including the enzymatic hydrolysis of extracellular ATP to form AMP, which is further metabolized to adenosine by ecto-5′-nucleotidase (CD73) (7). Adenosine regulates many physiological responses via activation of adenosine receptors (8), and it can also be taken up inside the cell via specific transporters (9) and phosphorylated back to AMP by adenosine kinase (10). A major function of adenosine is to reduce metabolic demand and conserve energy, but it is not known what role CD73 plays in this process, despite being the major extracellular adenosine regulator across different tissues and cell types (11, 12). Insights into CD73 mechanisms have particular relevance to liver biology and disease, since the mRNA expression and enzymatic activity of CD73 are significantly downregulated in human NAFLD, cirrhosis and HCC via transcriptional (13), post-transcriptional (14), and post-translational (15) mechanisms.

Although CD73 has ubiquitous expression(12), recent single cell profiling experiments show that its mRNA is zonally distributed in epithelial tissues, including the liver (16), intestine (17), and kidney (18). CD73 is transcriptionally induced by hypoxia (19) and CD73-generated adenosine directly protects multiple epithelial tissues, including the liver, against hypoxic injury (20-22). While previous studies demonstrated an allostatic function for CD73 in response to severe oxygen deprivation, presently it is not known if CD73 functions in the long-term maintenance of liver homeostasis. Addressing this question is important because oxygen tension across the hepatic lobule is a key determinant of physiological metabolic zonation (23). Metabolic zonation refers to the heterogeneous distribution of enzyme activities, resulting in periportal predominance of protein secretion, cholesterol synthesis, fatty acid oxidation, and ureagenesis, and pericentral predominance of glycolysis, lipogenesis and bile acid synthesis. This structured division of labor among healthy hepatocytes is disrupted in chronic liver diseases, such as NAFLD (24).

Given the above observations, and the known functions of adenosine in regulating hepatic glucose and lipid metabolism (25), we hypothesized that CD73 has non-redundant homeostatic functions in the liver. To test the hypothesis, we generated mice with a targeted deletion of the CD73-encoding gene *Nt5e* in hepatocytes and characterized their liver phenotypes using multiple approaches. Our findings reveal unanticipated age- and sex-dependent functions of CD73 in the long-term maintenance of hepatocyte metabolism and liver homeostasis. As such, these results add cellular-level understanding of this key enzyme, in particular its relatively underappreciated functions in epithelial tissues. Importantly, these findings have translational implications for human liver diseases, as well as anti-CD73 antibodies and inhibitors, which are currently undergoing clinical development for cancer (26) and COVID-19 immunotherapy (27).

## Experimental Procedures

Complete details of all reagents and experimental methods are provided in the Supplement.

### Mice

*Nt5e* floxed (*Nt5e*^fl/fl^) mice, generated as described in the Supplement, were bred with mice expressing Cre recombinase driven by the albumin promoter (both on C57BL/6J background) to generate the hepatocyte-specific CD73 knockout mice (CD73-LKO). WT and CD73-LKO littermates were fed normal chow diet and were co-housed (4-5/cage) in the pathogen-free animal facility at the University of North Carolina at Chapel Hill. Liver tissue and serum were collected and analyzed from male and female mice at 7 weeks, and between 21-42 weeks of age. All animals received humane care and experiments were approved by the Institutional Animal Care and Use Committee at UNC-Chapel Hill.

### Primary Hepatocyte Isolation and Treatment

Hepatocytes were isolated from 5-month-old mice by perfusion and collagenase digestion as described in the Supplement. Cells were treated 24 hours post-isolation with soluble human recombinant CD73 protein, AMP substrate, and a selective equilibrative nucleoside transporter I (ENT1) inhibitor, as specified in the figure legends. Proteomic analysis by liquid chromatography with tandem mass spectrometry (LC-MS/MS) was performed on freshly isolated WT and CD73-LKO hepatocytes and pathway analysis performed using the Ingenuity Pathway Analysis software.

### Statistical Analysis

Data were analyzed using GraphPad Prism and statistical significance determined by unpaired *t-*test or ANOVA. Data are expressed relative to WT or untreated controls. Error bars from all graphs indicate standard deviation (s.d.) for n≥3 samples and/or independent experiments, as specified in the figure legends. Significant outliers were tested based on Grubb’s test (α=0.05). The number of samples or independent experiments is indicated in the figure legends.

## Results

### Pericentral hepatocytes account for the bulk of CD73 expressed in the normal mouse liver

Using single cell RNA transcriptomic data from the Tabula Muris project (28), we compared *Nt5e* expression across different cell types in the mouse liver. Both FACS-sorted and microfluidic droplet-captured cells showed that hepatocytes are the primary cell types expressing *Nt5e* (Fig. 1A-D). Approximately 30% of hepatocytes, 7% of leukocytes and natural killer (NK) cells, and <2% of Kuppfer cells and liver sinusoidal endothelial cells (LSECs) express *Nt5e*, while cholangiocytes and B cells lack *Nt5e* expression (Fig. 1A-D). To determine if *Nt5e* presence correlates with CD73 protein, we performed co-immunofluorescence staining for CD73 and markers of hepatocytes, cholangiocytes, endothelial cells and Kupffer cells (Fig.1E). In agreement with the transcriptomic data, CD73 is present on hepatocytes, but absent from cholangiocytes, which are marked by keratin (K) K8/18 and K19 staining, respectively (Fig. 1E). In addition, subsets of endothelial cells, expressing CD31, and Kupffer cells, expressing F4/80, co-stained for CD73 (Fig.1E). However, the most abundant expression of CD73 was detected on the bile canalicular membranes of hepatocytes, as shown by co-staining with the tight junction marker protein ZO-1 (Fig.1F). Histological analysis also revealed that CD73 expression and enzymatic AMPase activity exhibited zonal distribution and were primarily associated with pericentral hepatocytes (Fig. 1G-H). The restricted zonal distribution of *Nt5e*/CD73 on hepatocytes suggests that it may serve potential specialized functions in liver homeostasis.

**FIG. 1.**
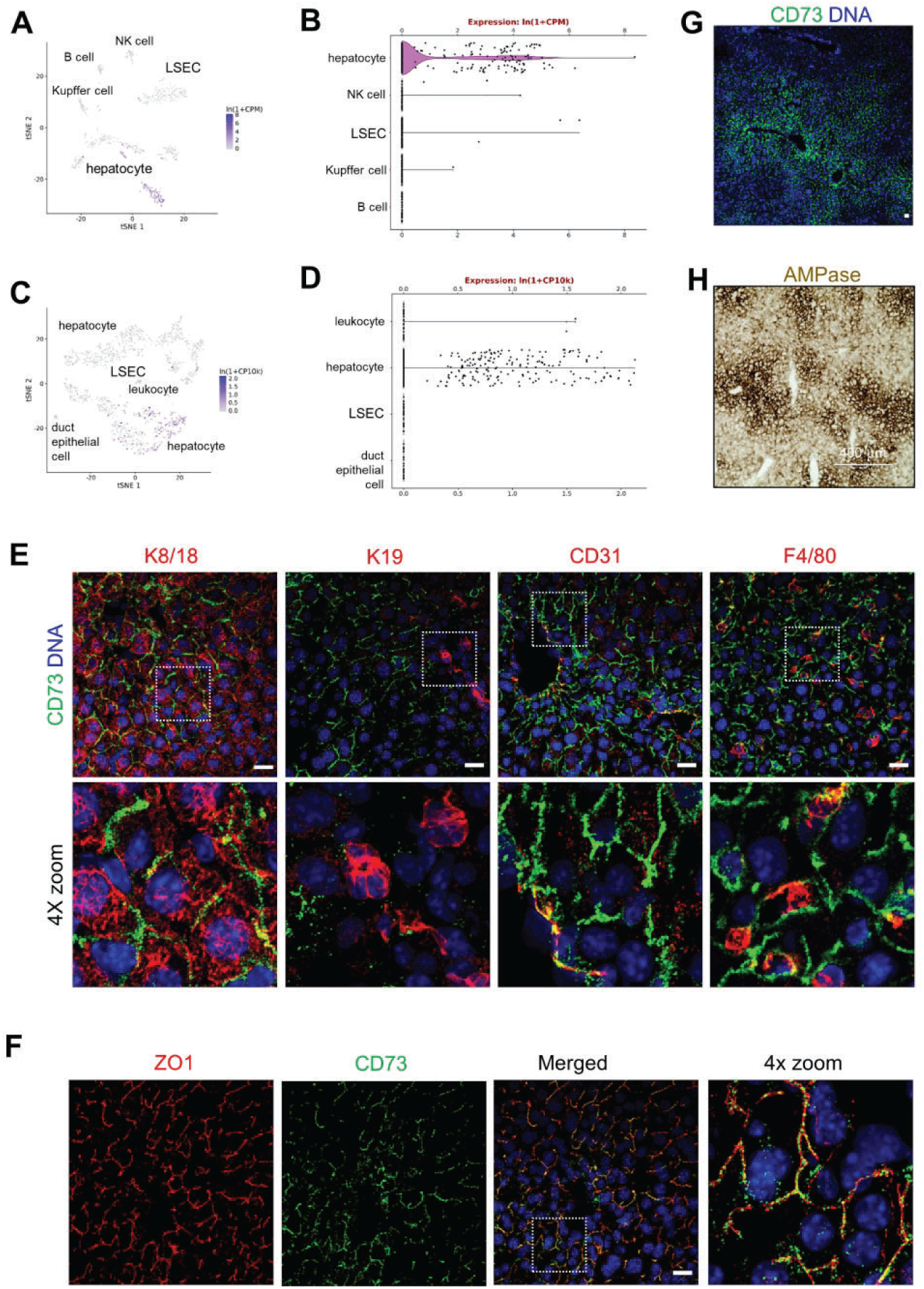
CD73 is primarily expressed on hepatocytes in normal mouse liver. tSNE and violin plots showing RNA-seq analysis of the mouse liver from **(A-B)** FACS-sorted and **(C-D)** microfluidic-droplet captured cells revealing highest expression of the mouse CD73-encoding gene *Nt5e* in hepatocytes. Data were obtained from Tabula Muris. **(E)** Fresh-frozen liver sections were stained with antibodies against CD73 (green), and the cell-specific markers keratins K8/18 (hepatocyte), K19 (cholangiocyte), CD31 (endothelium), F4/80 (Kupffer cells) in red. Bottom panels are magnified views of the boxed area in the top panels. DAPI-stained nuclei. Scale bar = 20μm. **(F)** Immunofluorescence staining for CD73 (green) in frozen liver sections showing co-localization with the bile-canalicular marker ZO1 (red). Right panel shows 4x magnified view of the boxed area in the merged panel. DAPI-stained nuclei. Scale bar=20μm. **(G)** AMPase activity in the mouse liver indicated by dark brown deposits in the presence of AMP substrate. Scale bar=400μm.

### Liver-specific CD73 knockout (CD73-LKO) mice develop normally and do not exhibit major liver abnormalities at a young age

To address the function of CD73 in the mammalian liver, we generated mice with a targeted deletion of the *Nt5e* gene in hepatocytes. The critical exon 2 of *Nt5e* was flanked by loxP sites, and deleted in the presence of Cre recombinase driven by the albumin (*Alb*) promoter. As expected, *Nt5e* was selectively targeted in the liver (Fig. 2A) and CD73 protein was absent from primary hepatocytes isolated from the CD73-LKO mice (Fig. 2B). Immunoblotting of total liver lysates revealed that CD73 levels were reduced by 75% and 67% in male and female CD73-LKO mice, respectively, compared to age-matched wild-type (WT) mice (Fig.2C-D). The expression of a functionally related GPI-anchored protein, tissue non-specific alkaline phosphatase (TNAP), was unaffected (Fig. 2C). Immunofluorescence analysis of liver tissues showed absence of CD73 from the apical membrane of hepatocytes in CD73-LKO mice (Fig. 2E) and this corresponded to significantly diminished *ecto*-AMPase activity in the pericentral region of CD73-LKO livers (Fig. 2F). Therefore, the CD73-LKO mice lack hepatocyte CD73 expression and are deficient in CD73-associated ecto-AMPase function.

**FIG. 2.**
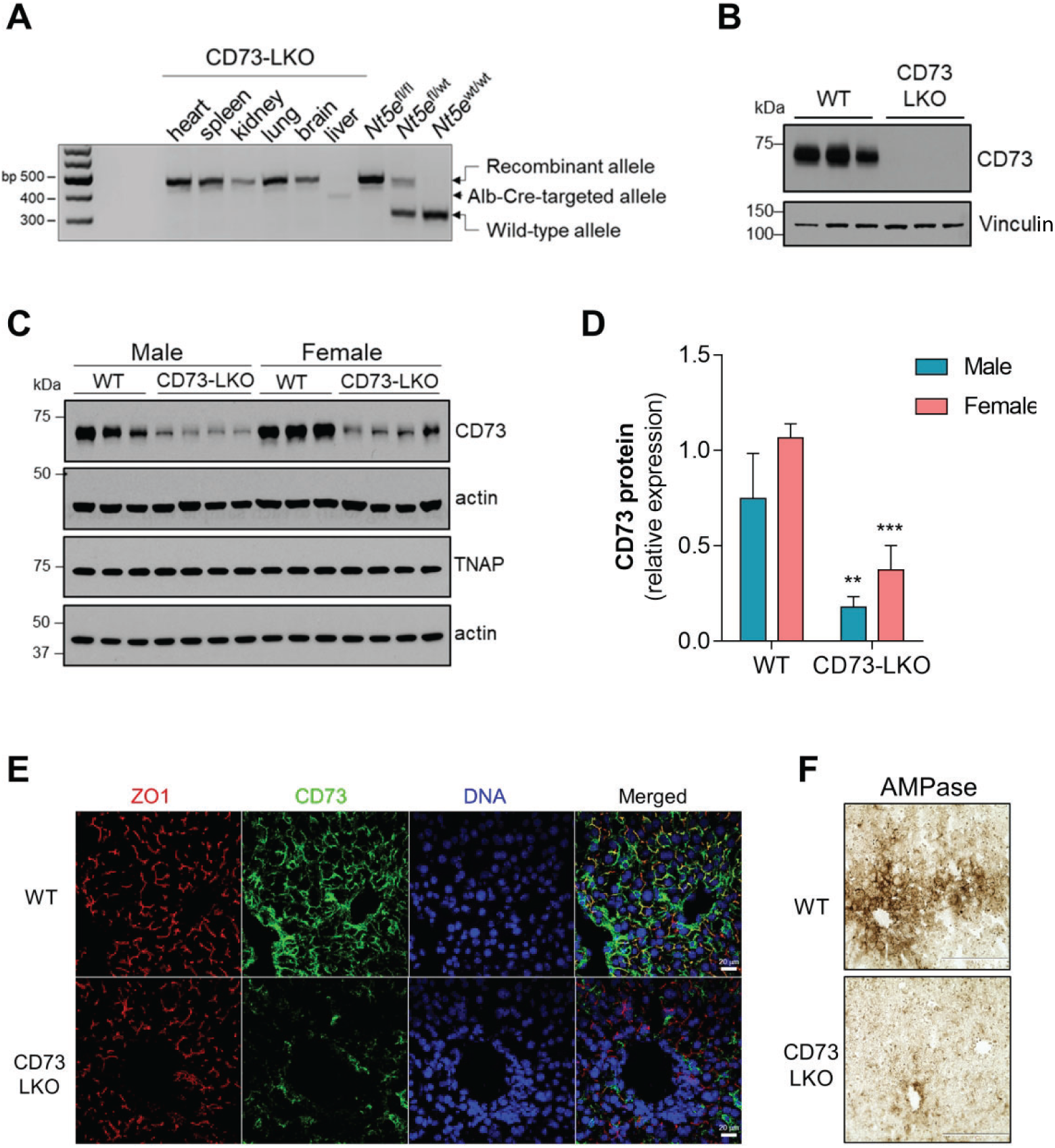
Generation of liver-specific CD73 knockout (CD73-LKO) mice. **(A)** PCR analysis of *Nt5e* generated a 349-bp fragment from the Cre-targeted allele in the CD73-LKO mouse liver. *Nt5e*^(wt/wt)^, *Nt5e*^(fl/wt)^, and *Nt5e*^(fl/fl)^ mice are controls. Representative immunoblots of CD73 in **(B)** primary hepatocytes, and **(C)** in total liver lysates from male and female WT and CD73-LKO mice. Actin, vinculin and TNAP immunoblots serve as controls. **(D)** Semi-quantitative analysis of CD73 protein expression based on band intensities in (C). ***p<0.01, ***p<0.001, two-way ANOVA. Error bars represent s.d. **(E)** Immunofluorescence staining for CD73 (green), tight junction protein ZO1 (red), and DAPI-stained nucleus (blue) on frozen liver tissue sections from WT (top) and CD73-LKO (bottom) mice. Scale bar=20μm. **(F)** AMPase activity in WT and CD73-LKO mice using formalin-fixed liver tissue sections. The brown deposits indicate ecto-AMPase activity in the presence of AMP substrate. Scale bar=400 μm.

To determine how this genetic manipulation affected liver function, we analyzed mice at different ages. Histological examination of liver tissues from 7-week old WT and CD73-LKO male mice did not reveal major differences (Fig.3A). In agreement with previous observations using constitutive *Nt5e*-knockout mice (22), the CD73-LKO mice did not show major defects in viability, growth, and development up to the mature adult stage at 21 weeks (Fig.3B). Both male and female CD73-LKO mice gained weight at the same rate (Fig. 3B), and had similar liver weight-to-body weight ratios compared to their WT counterparts (Fig. 3C). Taken together, these data show that hepatocyte CD73 is not required for normal mouse liver development and maturation.

**FIG. 3.**
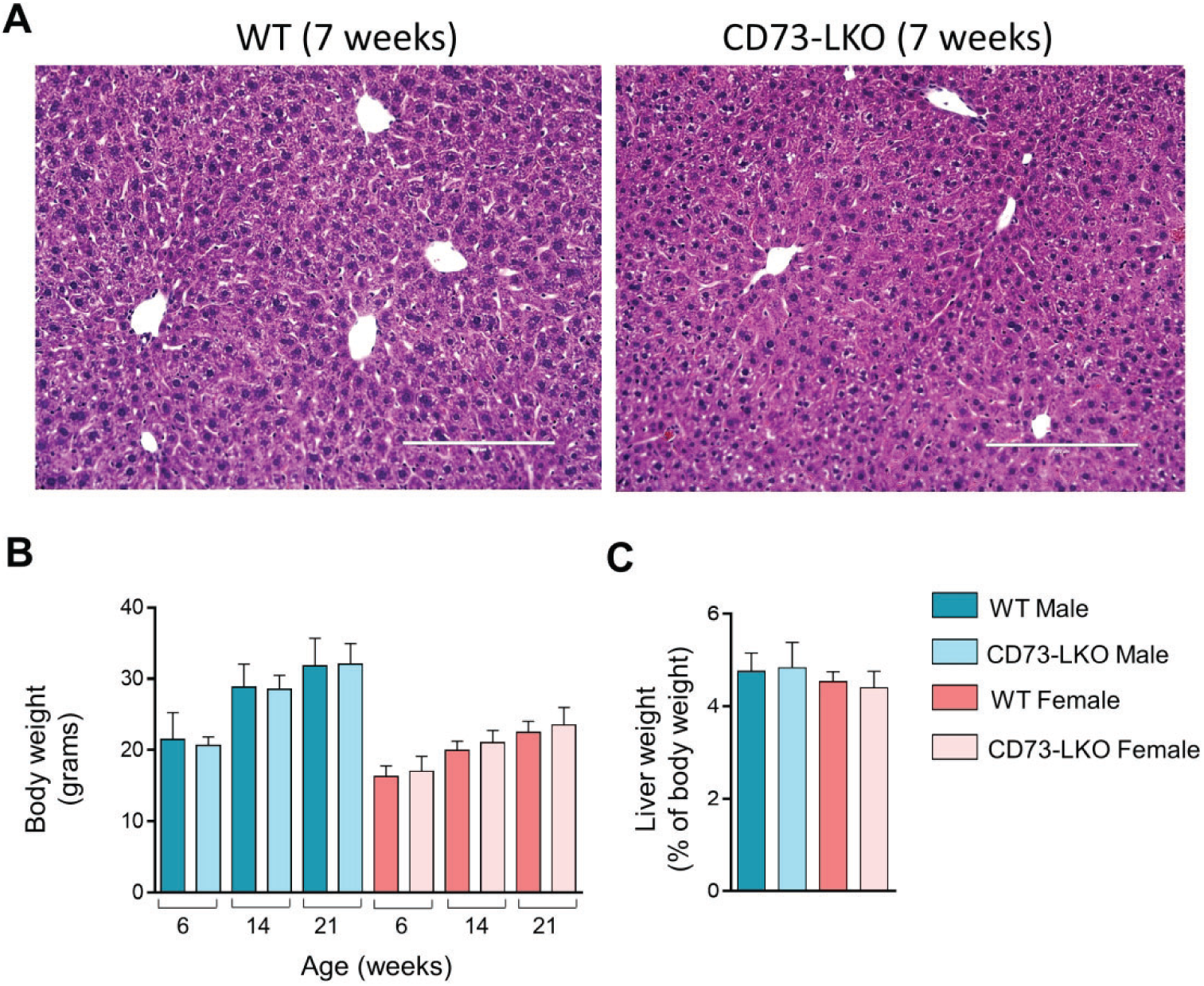
CD73-LKO mice develop normally and do not exhibit major liver abnormalities up to 21 weeks of age. **(A)** Formalin-fixed liver sections of 7-week old male mice stained with hematoxylin and eosin (H&E). Scale bar=200 μm. **(B)** Comparison of body weight from WT and CD73-LKO mice at 6-21 weeks of age. n=15-21 mice/group. **(C)** Analysis of liver-to-body weight ratios of 21-week old mice. Males: n=18 WT; n=21 CD73-LKO. Females: n=9 WT female, n=15 CD73-LKO. Error bars represent s.d.

### CD73-LKO mice develop spontaneous hepatocyte degeneration in a sex- and age-dependent manner

The zonal expression pattern of CD73 on pericentral hepatocytes prompted us to investigate the long-term consequences of CD73 deficiency, because pericentral hepatocytes are important for the homeostatic renewal in the mouse liver (29, 30). Therefore, we studied the liver phenotypes of standard chow-fed WT and CD73-LKO mice between 22-42 weeks (5-10 months) of age, which represents the period between the mature adult and middle-aged stages. Serum analysis showed a significant increase in ALT levels to 163.2±40.86 U/L in a mixed cohort of male and female CD73-LKO mice, compared to 65.73±18.27 U/L in WT mice (p<0.05) at 21-22 weeks of age (Fig. 4A). Histologically, we observed significant pericentral hepatocyte injury in 22-week old male CD73-LKO mice, which was not present in the female mice, or the corresponding age- and sex-matched WT controls (Fig. 4B). The hepatocyte injury and tissue damage were more extensive in older (42 week) compared to younger (22 week) male mice (Fig.4C), pointing to a progressive and age-dependent liver injury. Blinded quantitative histological analysis further unmasked male-predominant hepatocellular damage (p<0.05), which was defined by swelling and ballooning hepatocyte degeneration (Fig.4D). Another notable sex difference was that hepatocytes in female CD73-LKO appeared smaller (Fig.4B), which corresponded with decreased serum albumin levels (Fig. 4E). On the other hand, serum albumin was unaffected in the male CD73-LKO mice (Fig. 4E), pointing to potential metabolic compensation in male CD73-LKO hepatocytes. Furthermore, we observed a significant increase in serum blood urea nitrogen (BUN) levels in the male, but not female, CD73-LKO mice (Fig. 4F).

**FIG. 4.**
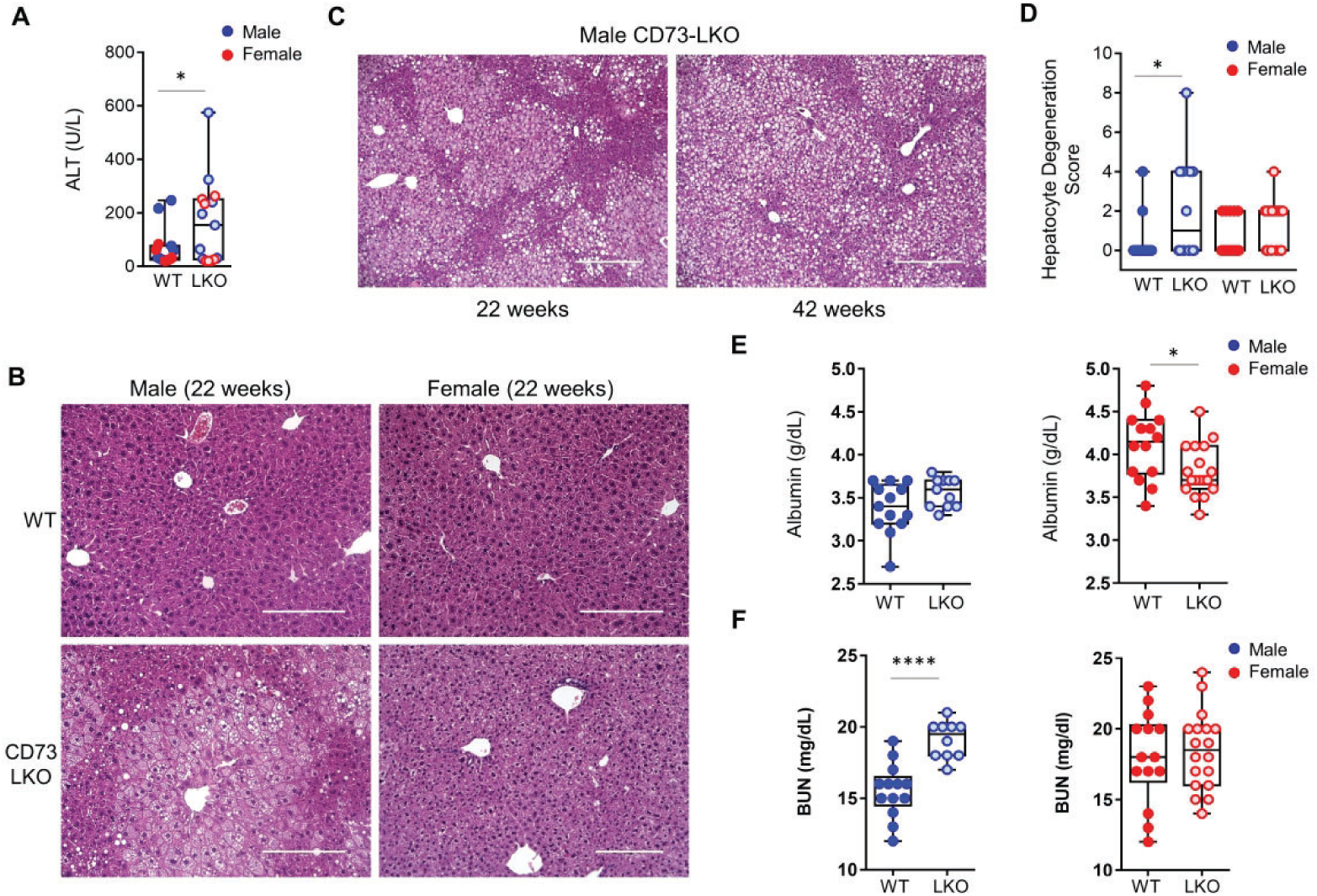
CD73-LKO mice develop spontaneous liver injury after 21 weeks of age in a sex-dependent manner. **(A)** Serum analysis of alanine aminotransferase (ALT) in 21-22 week old male (blue) and female (red) WT (filled circles) and CD73-LKO mice (open circles). Males: n=8 WT n=8 CD73-LKO; Females: n=7 WT n=7 CD73-LKO. *p<0.05, unpaired *t*-test. **(B)** Representative H&E images of formalin-fixed liver sections from 22-week old male and female WT and CD73-LKO mice. Scale bars=200 μm. **(C)** Representative H&E stained sections from 22-and 42-week old male CD73-LKO mice. Scale bars=200μm. **(D)** Quantification of blinded histological scoring of hepatocyte degeneration (defined as swelling and ballooning) in 21-42 week old mice. Scoring: 0=none, 4=mild swelling/focal ballooning, 8=severe swelling/extensive ballooning. (scores were averaged for animals with ballooning + swelling). *p<0.05; one-way ANOVA. Box-and-whiskers plots of serum albumin **(E)** and blood urea nitrogen (BUN) **(F)** in male (blue) and female (red) WT (filled circles) and CD73-LKO mice (open circles). Same animals as in panel D. Males: n=13 WT n=12 CD73-LKO; Females: n=14 WT n=18 CD73-LKO mice. *p<0.05, ****p<0.0001; unpaired *t*-test.

### Deficiency of hepatocyte CD73 leads to AMPK hypo-activation and perturbs metabolic homeostasis

In order to understand the cellular basis for the male-predominant histopathological changes, we performed proteomic profiling on freshly isolated hepatocytes from 5-month old male WT and CD73-LKO mice (Fig 5A). This analysis revealed upregulation in protein synthesis pathways associated with the transcriptional regulator EIF2, apoptosis, and nucleotide excision repair (NER). Pathways involved in fatty acid oxidation (PPARα/RXRα), cholesterol and stearate biosynthesis were also upregulated (Fig. 5A). On the other hand, hormone signaling (estrogen and aldosterone), amino acid metabolism (tRNA charging), nutrient sensing (mTOR, p70S6K), and overall cellular energy homeostasis (oxidative phosphorylation and AMPK) pathways were downregulated in the CD73-null hepatocytes (Fig. 5A). Consistent with abnormal lipid metabolism, as suggested by the proteomic analysis, male CD73-LKO mice developed significantly more micro- and macro-vesicular steatosis, compared to female mice (Fig. 5B-C).

**FIG. 5.**
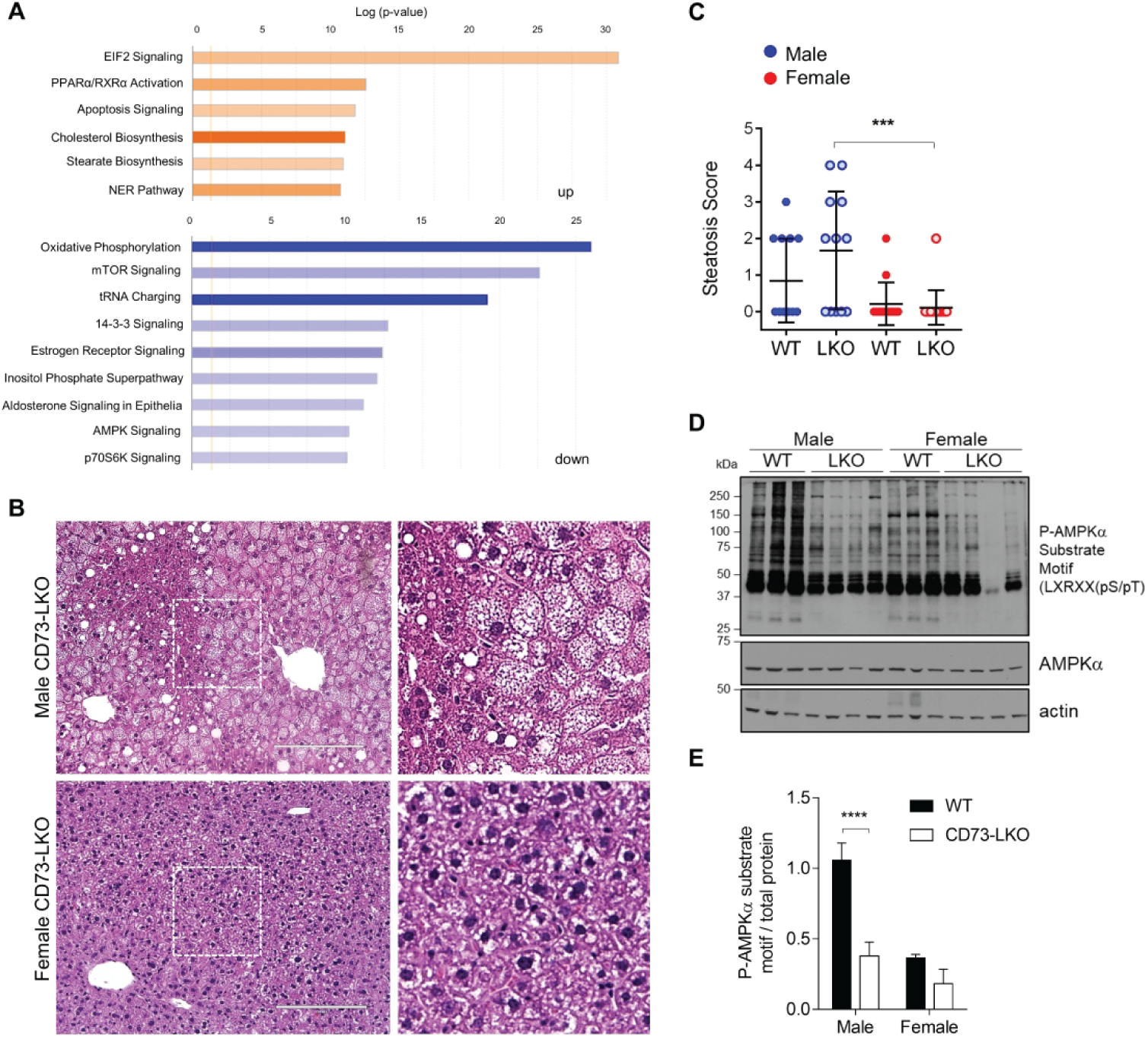
Metabolic imbalance and AMPK dysfunction in male CD73-LKO mouse hepatocytes and livers. **(A)** Ingenuity pathway analysis of proteomic results showing activated (up; orange) and inhibited (down; blue) pathways in freshly-isolated hepatocytes from male CD73-LKO mice relative to WT controls. **(B)** H&E staining of formalin-fixed liver sections. Magnified view of boxed areas in the left panel comparing the presence of steatosis in male and female CD73-LKO mice. Scale bar=200μm. **(C)** Quantification of blinded histological scoring of micro- and macro-vesicular steatosis. Scoring: 0=none, 1=mild; 2=moderate; 3=severe. (scores were averaged for animals with micro+macro steatosis). Males: n=13 WT; n=12 CD73-LKO; Females: n=14 WT n=18 CD73-LKO. ***p<0.001, one-way ANOVA. **(D)** Representative immunoblots of AMPK substrate phosphorylation in liver lysates from male and female mice. **(E)** Semi-quantitative analysis of AMPK substrate phosphorylation relative to total protein based on immunoblots in (D). ****p<0.0001, two-way ANOVA. Error bars represent s.d.

AMPK is a master metabolic regulator that acts as an ATP sensor and regulates glucose and lipid homeostasis. To determine if the metabolic dysregulation in the absence of CD73 was linked to altered AMPK signaling, we probed for AMPK substrate phosphorylation using a specific motif antibody [LXRXX (pS/pT)] (31). Despite equal expression of total AMPKα, male CD73-LKO mice had a dramatic reduction in the phosphorylation of AMPK substrates (Fig. 5D-E). On the other hand, baseline AMPK substrate phosphorylation was lower in WT female than male mice, and not changed significantly by CD73 deletion (Fig. 5D-E). The downregulation of AMPK substrate phosphorylation was not related to major changes in the abundance or subcellular distribution of the kinase (Fig. 6A), leading us to hypothesize that adenosine generated by CD73 promotes AMPK activity. In support of this hypothesis, extracellular addition of AMP significantly, and dose-dependently, induced AMPK activation (phosphorylation at Thr-172) in primary hepatocytes co-treated with soluble, enzymatically-active CD73 (Fig. 6B-C). To determine if this effect was dependent on the uptake of extracellular adenosine, we tested AMPK activation in the absence and presence of nitrobenzylthioinosine (NBTI), an inhibitor of the ENT1 adenosine re-uptake transporter (32). Although AMPK phosphorylation appeared blunted in the presence of (NBTI), this was not statistically-significant (Fig. 6D-E), suggesting multiple potential mechanisms. Combined, these data show that CD73, in the presence of extracellular AMP, activates AMPK in hepatocytes potentially via transport-dependent and -independent mechanisms.

**FIG. 6.**
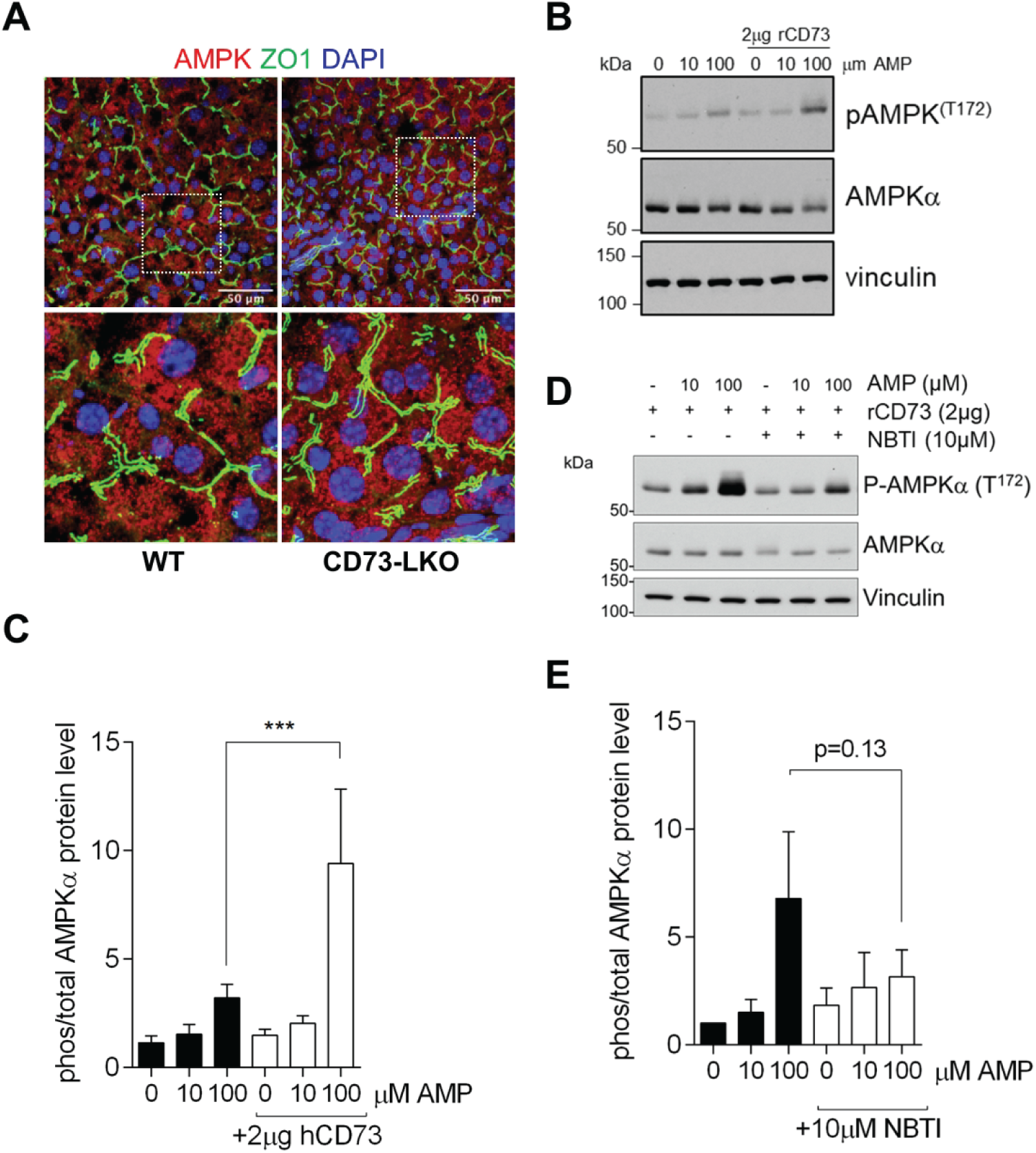
CD73-generated extracellular adenosine activates AMPK in primary mouse hepatocytes. **(A)** Fresh-frozen liver sections stained with antibodies against AMPK (red) and ZO1 (green) show similar AMPK distribution in WT and CD73-LKO mice. Bottom panels are magnified view of the boxed areas. **(B)** Isolated WT hepatocytes were treated with AMP and recombinant soluble CD73 (rCD73) at indicated concentrations. Representative immunoblots of total and phosho-AMPKα. **(C)** Quantification of phosphorylated/total AMPK. n=4 replicates. *p<0.001; two-way ANOVA. **(D)** Representative immunoblot analysis showing AMPK phosphorylation in WT hepatocytes treated with AMP, rCD73, and the adenosine transport inhibitor NBTI. **(E)** Quantification of phosphorylated/total AMPK in rCD73-treated hepatocytes, in the absence/presence of AMP and NBTI. n=3 replicates. two-way ANOVA. Error bars represent s.d.

### Mature adult CD73-LKO mice develop liver inflammation with male predominance in severity

Consistent with the known roles of adenosine (33) and AMPK (34) as anti-inflammatory mediators, in the CD73-LKO livers we observed induction in the pro-inflammatory cytokines IL-1β and TNFα, with the latter reaching statistical significance only in the female mice (Fig.7A). Blinded histological analysis showed presence of portal and lobular inflammation in both male and female CD73-LKO mice, but the male mice were more severely affected (Fig.7B-C). Profiling of CD73-associated adenosine pathway components, including the AMP substrate-generating enzyme CD39 (*Entpd1*) and adenosine receptors A1, A2A, A2B, and A3 (*Adora 1, 2A, 2B, and 3*), showed female-specific compensatory induction of these targets in the absence of hepatocyte CD73 (Fig.7D). These results demonstrate that loss of hepatocyte CD73 leads to spontaneous liver inflammation, and that in female mice this may be partly opposed by compensatory upregulation of anti-inflammatory adenosine signaling mediators to limit tissue damage.

**FIG. 7.**
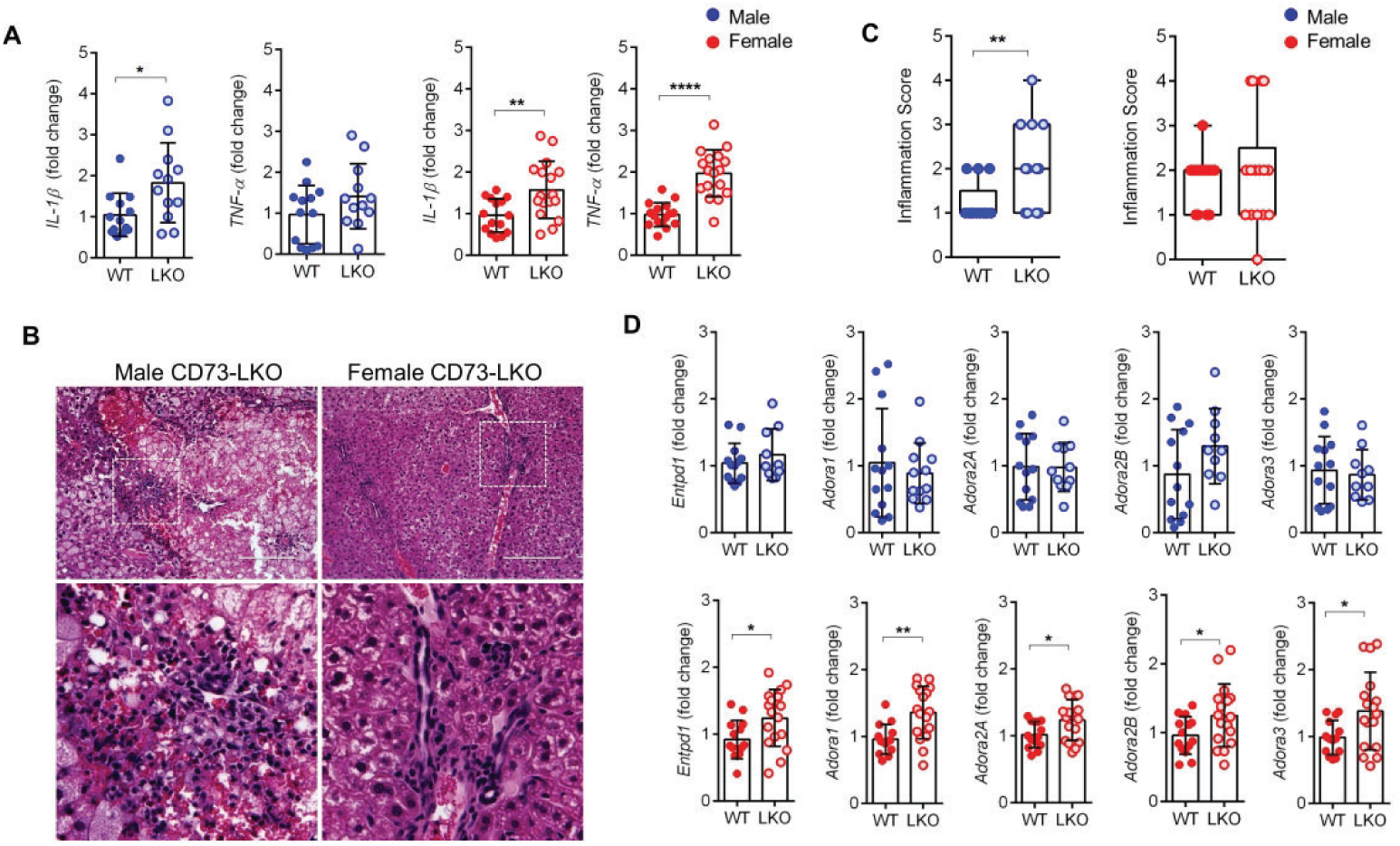
Loss of hepatocyte CD73 is associated with spontaneous liver inflammation. **(A)** Total liver mRNA analysis of pro-inflammatory markers IL1β and TNFα in male (blue) and female (red) mice aged 21-42 weeks. **(B)** Representative H&E image of inflammatory lesions in livers from 22-week old CD73-LKO mice. Boxed areas in the top panels are magnified in the bottom panels. **(C)** Quantification of blinded histological analysis of portal and lobular inflammation (scores were averaged for animals with portal + lobular inflammation). Scoring: 0=none, 2=minimal; 4=mild; 6=moderate; 8=severe. **(D)** Gene expression analysis of ectonucleotidase CD39 (*Entpd1*) and adenosine receptors A1R, 2AR, 2BR, 3R (*Adora1, 2a, 2b, 3*) using total liver mRNA in male and female WT and LKO mice. n=13 WT male, 12 LKO male, 14 WT female, 18 LKO female. *p<0.05, **p<0.01, ****p<0.0001, unpaired *t*-test. Error bars represent s.d.

## Discussion

### Physiological hepatoprotection of CD73 and adenosine

Our study demonstrates the importance of CD73, a major extracellular AMPase, for the long-term metabolic integrity of the mammalian liver. Previous seminal work utilizing constitutive CD73 knockout mice established CD73 as a key factor in maintaining epithelial integrity via adenosine-mediated protection of the tissue barrier function, especially during hypoxia (19, 22, 35, 36), while our present findings provide new insights into the homeostatic mechanisms that are specifically attributed to CD73 on hepatocytes. The male-predominant, spontaneous development of liver disease that we observed in CD73 LKO mice points to a physiological protective mechanism of CD73 that exhibits a sexual dimorphism. We conclude that AMPK hypo-activation, likely secondary to extracellular adenosine deficiency, contributes to the liver injury in the CD73 LKO male mice. CD73 is significantly downregulated under conditions of severe and persistent chronic stress in the mouse liver (13), and in chronic human liver diseases of different etiologies (14, 15). Future studies addressing the detailed mechanisms of how CD73 supports metabolic homeostasis in hepatocytes throughout the lifespan, and in a sex-dependent manner, will have important implications for understanding mammalian liver biology as well as disease development and progression. To that end, the *Nt5e*^fl/fl^ and CD73-LKO mice are new tools for studying the cell-specific mechanisms of human liver diseases that are driven by perturbed metabolism and inflammation, such as NAFLD (37).

### CD73 as a mediator of sex-dependent liver injury responses

Biological sex and reproductive hormones exert a significant effect over individual variability to human disease development and progression (38). This is especially evident in chronic liver disease, which currently ranks among the top ten causes of death in men, but not in women (39). Both male sex and older age are associated with worse survival and greater incidence of HCC among patients with biopsy-confirmed NAFLD (40). However, the basic biological mechanisms of how age and biological sex affect patient susceptibility to disease development and progression have been difficult to understand, especially in the case of NAFLD (41). Our results presented here add new insight into how biological sex impacts metabolic stress related to altered purinergic signaling in the liver. CD73 represents a potential target for extending liver function and preventing disease progression.

### Cross-talk between CD73 and AMPK in hepatocytes

CD73 is strongly induced in epithelial tissues under hypoxic conditions via hypoxia-inducible factor-1 (HIF-1) (19). Rapid and sustained CD73 upregulation during hepatic ischemia confers adenosine-dependent hepatoprotection (21). The latter is very similar to the reported hepatoprotective effect of AMPK during ischemic preconditioning (42), although a functional link between these pathways has not been established. At 5 months of age, AMPK activity in mouse liver is significantly augmented by hypoxia, but this regulation is lost at older age (43). The age-dependent activation of AMPK by hypoxia in the liver coincides with the age of onset of spontaneous liver injury in the CD73 LKO mice, which became apparent at 5 months in our study animals. Based on this, we propose that the detrimental effects stemming from CD73 deficiency in hepatocytes may be associated with impaired capacity of the mature liver to calibrate oxygen responses properly via AMPK. *In vitro* work in cell lines, and *in vivo* studies using lower organisms, have linked AMPK and HIF-1 cross-talk with cellular homeostasis and organismal longevity through their roles in balancing intracellular reactive oxygen species (ROS) (44, 45). The purinergic signaling pathway also plays important roles in oxidative stress responses in the liver and is disrupted in liver disease (46). As a key regulator of purinergic signaling, and the major extracellular enzymatic source of the immuno-suppressive and anti-inflammatory mediator adenosine, CD73 represents a promising target for metabolic inflammation, which is considered a major driver of chronic human diseases and associated comorbidities (37, 47).

### Anti-inflammatory effects of adenosine in the liver

In various liver injury models, adenosine exerts protective anti-inflammatory effects through the activation of the G_s_-protein coupled adenosine receptor 2A (A_2A_R) on hepatocytes and immune cells (33, 48, 49). For example, genetic deletion or pharmacological inhibition of the A_2A_R causes severe inflammation and liver damage following concanavalin A (Con A)-induced liver injury in a mixed cohort of male and female mice (48). Whole-body hypoxia treatment provided protection against Con A-induced liver inflammation (49). Although the detailed mechanism behind the hepatoprotective effects of hypoxia in liver inflammation are not clear, it is plausible that hypoxia-mediated CD73 induction, followed by increased extracellular adenosine production, may be important during immune-mediated liver damage, which remains to be examined in detail in future studies. CD73 protein was shown to be upregulated in a high fat diet (HFD) model of liver injury performed in male mice, where the hepatic protein expression of CD39 and A_2A_R were also increased (33). The upregulation of these purinergic factors was likely a protective mechanism, because the absence of A_2A_R led to more severe inflammation, steatosis and impaired insulin sensitivity after HFD feeding (33).

The hypo-activation of AMPK and liver injury in the CD73 LKO mice that we observed are similar to the findings from another study employing a mixed cohort of older (12-16 months) male and female mice, which found that the absence of the A_2A_R receptor leads to hypo-activation of AMPK and metabolic dysregulation, specifically obesity, hyperglycemia and glucose intolerance (50). The relative protection that we observed in the female CD73 LKO mice in our study could be attributed to compensatory induction of adenosine receptors, in particular A_2A_R and A_2B_R. Future studies addressing the contribution of adenosine receptor-dependent vs –independent mechanisms and how these impact the sex-dependent physiological and pathophysiological hepatic responses in CD73 LKO mice remain to be investigated. Such studies should identify additional factors that sustain the long-term metabolic and regenerative capacity of the liver, and uncover new strategies for harnessing the potent activity of adenosine to achieve hepatoprotection.

## List of Abbreviations

AMP: adenosine monophosphate
AMPK: AMP-activated protein kinase
AR: adenosine receptor
ATP: adenosine triphosphate
CD73: ecto-5′-nucleotidase
CD73-LKO: liver-specific CD73 knockout
HCC: hepatocellular carcinoma
MAFLD: Metabolic-Associated Fatty Liver Disease
NAFLD: non-alcoholic fatty liver disease
NT5E: CD73-encoding gene
TNAP: tissue non-specific alkaline phosphatase

## Supplementary File 1. Detailed Experimental Procedures

### Chemicals and Reagents

The following chemicals and reagents were purchased from Sigma-Aldrich (St. Louis, MO): bovine serum albumin (A2153), Adenosine 5′-(α,β-methylene) diphosphate (M3763), insulin (I0516), glucagon (G2044), hydrocortisone (H0888), levamisole (196142), calcium chloride dehydrate (223506), glucose (G8270), Percoll (P1644), and EGTA (E3889). 4X Laemmli Sample buffer (1610747) was purchased from BioRad Laboratories (Hercules, CA).

### Generation of hepatocyte-specific CD73 KO mice

CD73 floxed (CD73^fl/fl^) mice were generated by Cyagen Bioscience, Inc. (Santa Clara, CA). Briefly, exon 2 of the CD73-encoding gene, *Nt5e* (GenBank accession number: NM_011851.4, Ensembl: ENSMUSG00000032420), was targeted to generate a frameshift mutation from downstream exons. BAC clone RP23-137M4 from the C57BL/6J library was the template for the targeting vector that contains a Neo cassette flanked by Frt sites, and *Nt5e* exon 2 region flanked by LoxP sites. Linearized targeting vector was electroporated into embryonic stem (ES) cells, followed by Neomycin selection of resistant clones. Targeted ES clones were injected into blastocysts and transferred into surrogate mothers to obtain chimera lines. The conditional knockout allele was obtained after Flp-mediated recombination. Heterozygote (flox/wt) mice were bred with each other to obtain flox/flox. For this study, CD73^fl/fl^ mice were backcrossed to C57BL/6J WT mice four times. To generate the CD73 hepatocyte-specific knockout mice (CD73-LKO), CD73^fl/fl^ mice were bred with Cre mice driven by the albumin promoter (Stock 003574, The Jackson Laboratory) (1). Genotyping was performed at 7-10 days of age. WT and LKO littermates were fed normal chow diet and weaned at 21 days.

### Genotyping

PCR analysis using DNA extracted from toe clips and DreamTaq Green PCR Master Mix 2X (Thermo Scientific, #K1089) was performed to identify WT and CD73-LKO mice. Flox PCR analysis used primers Fl-F2 (5’-AGCACATTTAGTTTGAAATCCC-3’), and Fl-R2 (5’-AAACAGACTTCTTGATTGGCAT-3’). To identify the presence of Cre recombinase, we used primers Fabpi-200R (5’-TAGAGCTTTGCCACATCACAGGTCATTCAG-3’), Fabpi-200F (5’-TGGACAGGACTGGACCTCTGCTTTCCTAGA-3’), Cre-281R (5’-TCGCCATCTTCCAGCAGG-3’), Cre-281F (5’-CCATCTGCCACCAGCCAG-3’). Amplicons were analyzed using 2% agarose gel electrophoresis. The presence of a 426-bp and 286-bp bands indicate floxed CD73 and WT allele, respectively. A 300-bp and 200-bp band is positive for Cre recombinase.

### Serum Analysis

Blood was collected by cardiac puncture from male and female WT and CD73-LKO mice between 5-9 months of age. Serum was analyzed for alanine aminotransferase (ALT), albumin, and blood urea nitrogen (BUN) using the VetScan VS2 Analyzer and VetScan mammalian liver profile reagent rotor (Abaxis, Union City, CA).

### Blinded histological analysis of liver sections

Liver histology in the mice was assessed by a board-certified pathologist (KGG) in a blinded fashion. Criteria for assessing inflammation, hepatocyte degeneration and steatosis are presented in the table below. For the quantification plots presented in the main figures, numerical scores were assigned based on the severity in each category. For each animal, scores were calculated for inflammation (average of portal and lobular); hepatocyte degeneration (average of swelling and ballooning) and steatosis (average of micro- and macrovesicular).

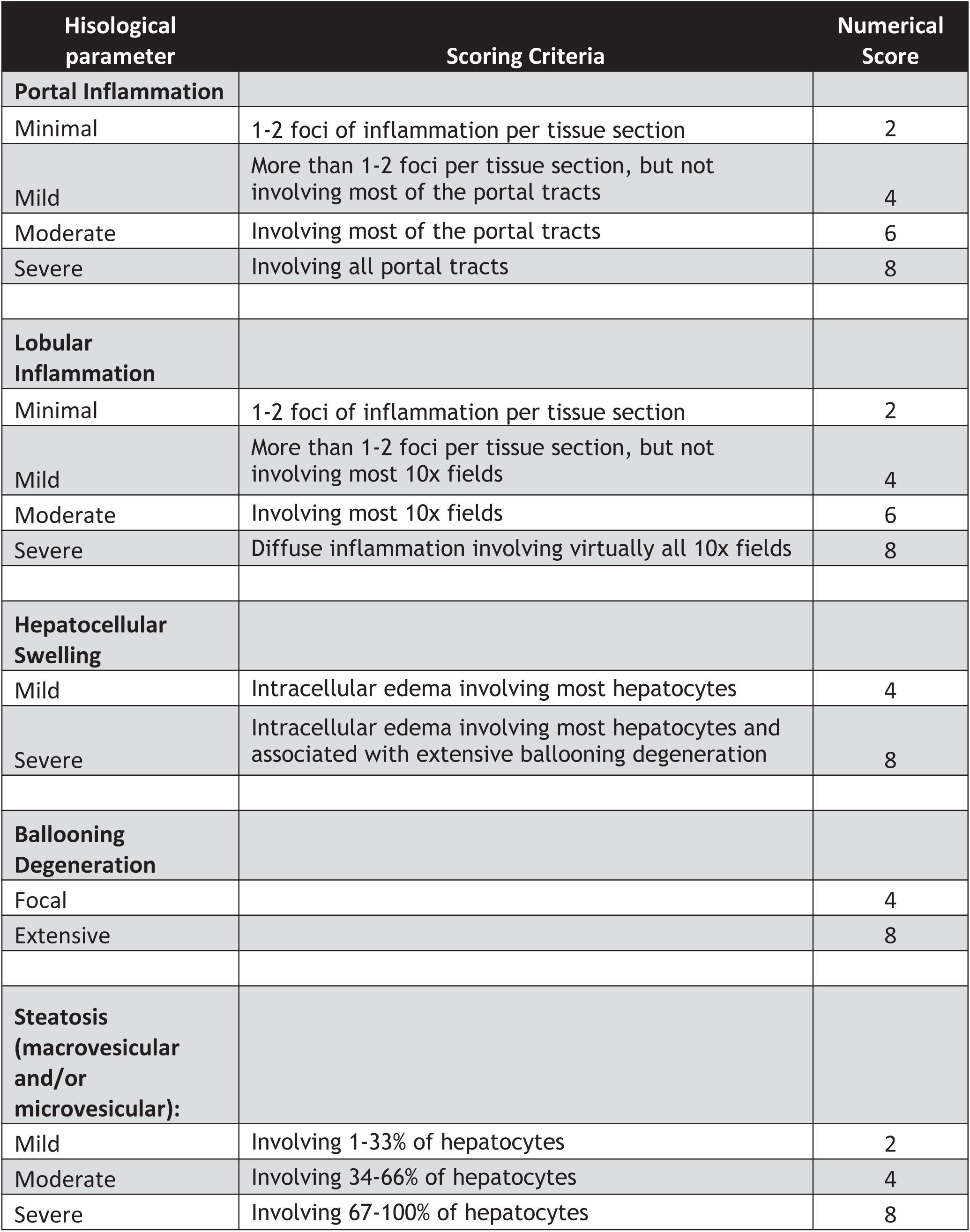

### Primary Hepatocyte Isolation and Treatment

Hepatocyte isolation was performed on 5-month-old CD73^fl/fl^ mice that weighed between 20-35g. Mice were anesthetized with Nembutal (40-60 mg/kg, IP) to achieve deep anesthesia. Portal vein was cannulated and liver was perfused with 50 ml of 37°C sterile buffer I solution (50 mM EGTA, 1M glucose, 1% pen/strep in calcium- and magnesium-free Hank’s Balanced Salt Solution (HBSS)) at a rate of 7 ml/min. Liver digestion was performed with 40 mL of 37°C sterile buffer II solution (1M CaCl_2_, 1M glucose, 1% pen/strep, and 3600 U of Collagenase IV in calcium- and magnesium-free Hank’s Balanced Salt Solution (HBSS) at a rate of 7 ml/min. Livers were surgically removed and hepatocytes were isolated in cold 1X DMEM/pen/strep media by size exclusion using 100 μm and 70 μm filters, respectively. Cells were subsequently washed at 120 x *g* for 5 min at 4°C. Live/dead cell exclusion was performed by Percoll gradient (1:1 Percoll/DMEM media) and centrifugation at 120 x *g* for 5 min at 4°C. Hepatocytes were cultured in complete hepatocyte medum (1X DMEM, 10% fetal bovine serum, 1% pen/strep, 0.02 mg/ml insulin, 0.0284 mg/ml glucagon, and 0.015 mg/ml hydrocortisone) at a density of 200,000 cells/well in 24-well collagen I-coated plates (Gibco #A1142802) or in 14-mm collagen I-coated glass bottom plates as previously published (2). Cells were treated 24 hours post-isolation with 2μg of active, soluble human recombinant CD73 protein and 0, 10, 100 μM AMP substrate for 30 minutes. S-(4-Nitrobenzyl)-6-thioinosine (NBTI, Sigma-Aldrich, #N2255) was used to selectively inhibit the equilibrative nucleoside transporter I (ENT1). Cells were collected and analyzed by immunoblot.

### Immunoblot

Total liver protein lysates were extracted from 25 mg of liver tissue from the left lobe or from cultured primary WT hepatocytes using ice-cold lysis buffer (50 mM n-octyl-β-D-glucopyranoside, 1X protease inhibitors (Roche, #05892970001), and 1X phosphatase inhibitors (Roche, #04906837001) in 1X PBS). Protein lysates were resolved in 10% SDS-PAGE gels, and transferred to a nitrocellulose membrane. After blocking in 5% milk/TBST for 1hr at room temperature (RT), the membranes were incubated with anti-CD73 (clone D7F9A, #13160), phospho-AMPKα (T172, clone 40H9, #2535), total AMPKα (clone D5A2, #5831), and phospho-AMPK substrate motif (LXRXX(pS/pT), #5759S) (Cell Signaling); pan actin (ACTN05, Thermo Fisher #MA5-11869), and anti-vinculin (clone hVIN-1, Sigma #V9131) in 5% BSA/TBST overnight at 4°C. Secondary antibodies (Sigma, #A4416, #A6154)(1:5,000-10,000) were incubated for 1 hour at RT in 5% milk/TBST. Gels were stained with Coomassie Blue, and membranes with Ponceau S stain.

### RNA and Quantitative PCR

Total RNA was isolated from the left lobe of the liver and was extracted according to the manufacturer’s protocol (RNeasy Mini Kit, Qiagen, #74104). A total of 2 mg RNA was converted to cDNA using the high capacity cDNA reverse transcription kit (Applied Biosystems, #4368813). PCR was performed using SYBR Green PCR master mix (Applied Biosystems, #A25742), and carried out in a Quantstudio™ 6 Flex System (Applied Biosystems). The delta CT was normalized to 18S, and was expressed in fold change. The primer sequences are: *Adora1* F: 5’-TGTGACCACCACCCAGAGTA -3’ R: 5’-GCAGAGACTGGGACAAGGAG -3’. *Adora2a* F: 5’-GAAGCAGATGGAGAGCCAAC -3’ R: 5’-GAGAGGATGATGGCCAGGTA -3’. *Adora2b* F: 5’-CCTTTGGCATTGGATTGACT -3’ R: 5’-AAAATGCCCACGATCATAGC -3’. *Adora3* F: 5’-TCATACCGGAAGGAATGAGC -3’ R: 5’-AGCTTGACCACCCAGATGAC -3’. *Entpd1* F: 5’-TACCACCCCATCTGGTCATT-3’ R: 5’-GGACGTTTTGTTTGGTTGGT -3’. *IL1β* F: 5’-TCGCTCAGGGTCACAAGAAA-3’ R: 5’-CATCAGAGGCAAGGAGGAAAAC-3’. *TNFα* (Mar. 2, 2018) F: 5’-AGGCTGCCCCGACTACGT-3’ R: 5’-GACTTTCTCCTGGTATGAGATAGCAAA -3’.

### Liver Tissue Staining

Liver tissues from the left lobe were fixed with paraformaldehyde and paraffin-embedded, or flash frozen using OCT media. Paraffin-embedded tissues were cut in 10mm sections and stained with hematoxylin and eosin. Images were analyzed at 20X objective using EVOS™ FL auto imaging system. Frozen tissues were cut in 10 μm sections and stained by immunofluorescence. Briefly, frozen tissue sections were fixed in cold 10% normal buffered formalin (Fisherbrand, #032-059), followed by cold acetone (Fischer-Scientific, #A19-1) for 10 minutes each. Tissues were blocked in 2% normal goat serum/2.5%BSA/1X PBS for 1 hour at RT. Primary antibodies CD73 (clone TY/23, BD Pharmingen #550738), K19-AF647 (Abcam #ab192980), K18/8 (clone 8592), ZO1 (clone D6L1E, Cell Signaling #13663) in blocking solution were incubated overnight at 4°C. Tissues were washed in 1X PBS, before adding secondary antibodies anti-rat 488 (Thermo Fisher, #a11006) and anti-rabbit 568 (Thermo Fisher, #a11079) for 1 hour at RT. The nuclei were stained using DAPI (Invitrogen, #D1306). Images were analyzed at 40X oil objective using Zeiss LSM 880 confocal microscope.

### CD73 Enzymatic Activity and Enzyme Histochemistry

Frozen liver tissues sections of 10μm thickness were fixed in 10% neutral buffered formalin for 5 min at 4°C. Enzyme histochemistry was performed using our published protocol (3). Images were analyzed at 20X objective using EVOS™ FL auto imaging system.

### Proteomic Analysis by Liquid Chromatography with Tandem Mass Spectrometry (LC-MS/MS)

Freshly isolated primary hepatocytes from 3 WT and 3 CD73-LKO mice were lysed using 8M urea. Protein lysates were digested with LysC and trypsin. Peptides samples were cleaned using C18 spin columns (Pierce), then peptide quantitation was conducted. A total of 350 mg per sample were dried and labeled using TMT 6-plex™ Isobaric Label Reagent Set (Thermo Fisher, #90061). Label efficiency was evaluated and found to be >99% for all TMT channels. Samples were mixed, then subjected to fractionation into 96 fractions by high-pH reversed-phase LC. Fractions were concatenated into 24 samples. Each sample (∼1mg) was analyzed by LC-MS/MS using the QExactive HF (Thermo Scientific) for a total of 25 analyses. Proteins were identified and quantified using MaxQuant utilizing both a reviewed (∼18,000 proteins) mouse database appended with a common contaminants database. Core analysis of LC-MS/MS data set were analyzed using Ingenuity Pathway Analysis (Qiagen) and filtered based on a log (*P* value of 1E^-10^) and an absolute z-score of 1.

### Statistical Analysis

Data were analyzed using unpaired *t-*test or two-way ANOVA built in GraphPad Prism. Data are presented relative to WT or untreated controls. Error bars from all graphs indicate standard deviation (s.d.) for n≥3 samples or independent experiments. *P* values are denoted within each respective figure panel. Outliers were tested based on Grubb’s test (α=0.05). Number of samples or independent experiments is indicated in the figure legends.

## References

1. Younossi Z, Anstee QM, Marietti M, Hardy T, Henry L, Eslam M, George J, et al. Global burden of NAFLD and NASH: trends, predictions, risk factors and prevention. Nat Rev Gastroenterol Hepatol 2018;15:11–20.

2. Anstee QM, Reeves HL, Kotsiliti E, Govaere O, Heikenwalder M. From NASH to HCC: current concepts and future challenges. Nat Rev Gastroenterol Hepatol 2019;16:411–428.

3. Cotter TG, Rinella M. Nonalcoholic Fatty Liver Disease 2020: The State of the Disease. Gastroenterology 2020;158:1851–1864.

4. Eslam M, Sanyal AJ, George J, International Consensus P. MAFLD: A Consensus-Driven Proposed Nomenclature for Metabolic Associated Fatty Liver Disease. Gastroenterology 2020;158:1999–2014 e1991.

5. Hyams DE, Isselbacher KJ. Prevention of Fatty Liver by Administration of Adenosine Triphosphate. Nature 1964;204:1196–1197.

6. Cortez-Pinto H, Chatham J, Chacko VP, Arnold C, Rashid A, Diehl AM. Alterations in liver ATP homeostasis in human nonalcoholic steatohepatitis: a pilot study. JAMA 1999;282:1659–1664.

7. Zimmermann H. 5’-Nucleotidase: molecular structure and functional aspects. Biochem J 1992;285 (Pt 2):345–365.

8. Borea PA, Gessi S, Merighi S, Vincenzi F, Varani K. Pharmacology of Adenosine Receptors: The State of the Art. Physiol Rev 2018;98:1591–1625.

9. Griffiths M, Beaumont N, Yao SY, Sundaram M, Boumah CE, Davies A, Kwong FY, et al. Cloning of a human nucleoside transporter implicated in the cellular uptake of adenosine and chemotherapeutic drugs. Nat Med 1997;3:89–93.

10. Spychala J, Datta NS, Takabayashi K, Datta M, Fox IH, Gribbin T, Mitchell BS. Cloning of human adenosine kinase cDNA: sequence similarity to microbial ribokinases and fructokinases. Proc Natl Acad Sci U S A 1996;93:1232–1237.

11. Colgan SP, Eltzschig HK, Eckle T, Thompson LF. Physiological roles for ecto-5’-nucleotidase (CD73). Purinergic Signal 2006;2:351–360.

12. Minor M, Alcedo KP, Battaglia RA, Snider NT. Cell type-and tissue-specific functions of ecto-5’-nucleotidase (CD73). Am J Physiol Cell Physiol 2019;317:C1079–C1092.

13. Snider NT, Griggs NW, Singla A, Moons DS, Weerasinghe SV, Lok AS, Ruan C, et al. CD73 (ecto-5’-nucleotidase) hepatocyte levels differ across mouse strains and contribute to mallory-denk body formation. Hepatology 2013;58:1790–1800.

14. Snider NT, Altshuler PJ, Wan S, Welling TH, Cavalcoli J, Omary MB. Alternative splicing of human NT5E in cirrhosis and hepatocellular carcinoma produces a negative regulator of ecto-5’-nucleotidase (CD73). Mol Biol Cell 2014;25:4024–4033.

15. Alcedo KP, Guerrero A, Basrur V, Fu D, Richardson ML, McLane JS, Tsou CC, et al. Tumor-Selective Altered Glycosylation and Functional Attenuation of CD73 in Human Hepatocellular Carcinoma. Hepatol Commun 2019;3:1400–1414.

16. Ben-Moshe S, Shapira Y, Moor AE, Manco R, Veg T, Bahar Halpern K, Itzkovitz S. Spatial sorting enables comprehensive characterization of liver zonation. Nat Metab 2019;1:899–911.

17. Moor AE, Harnik Y, Ben-Moshe S, Massasa EE, Rozenberg M, Eilam R, Bahar Halpern K, et al. Spatial Reconstruction of Single Enterocytes Uncovers Broad Zonation along the Intestinal Villus Axis. Cell 2018;175:1156–1167 e1115.

18. Broeker KAE, Fuchs MAA, Schrankl J, Kurt B, Nolan KA, Wenger RH, Kramann R, et al. Different subpopulations of kidney interstitial cells produce erythropoietin and factors supporting tissue oxygenation in response to hypoxia in vivo. Kidney Int 2020;98:918–931.

19. Synnestvedt K, Furuta GT, Comerford KM, Louis N, Karhausen J, Eltzschig HK, Hansen KR, et al. Ecto-5’-nucleotidase (CD73) regulation by hypoxia-inducible factor-1 mediates permeability changes in intestinal epithelia. J Clin Invest 2002;110:993–1002.

20. Tak E, Jung DH, Kim SH, Park GC, Jun DY, Lee J, Jung BH, et al. Protective role of hypoxia-inducible factor-1alpha-dependent CD39 and CD73 in fulminant acute liver failure. Toxicol Appl Pharmacol 2017;314:72–81.

21. Hart ML, Much C, Gorzolla IC, Schittenhelm J, Kloor D, Stahl GL, Eltzschig HK. Extracellular adenosine production by ecto-5’-nucleotidase protects during murine hepatic ischemic preconditioning. Gastroenterology 2008;135:1739–1750 e1733.

22. Thompson LF, Eltzschig HK, Ibla JC, Van De Wiele CJ, Resta R, Morote-Garcia JC, Colgan SP. Crucial role for ecto-5’-nucleotidase (CD73) in vascular leakage during hypoxia. J Exp Med 2004;200:1395–1405.

23. Gebhardt R. Metabolic zonation of the liver: regulation and implications for liver function. Pharmacol Ther 1992;53:275–354.

24. Hall Z, Bond NJ, Ashmore T, Sanders F, Ament Z, Wang X, Murray AJ, et al. Lipid zonation and phospholipid remodeling in nonalcoholic fatty liver disease. Hepatology 2017;65:1165–1180.

25. Koupenova M, Ravid K. Adenosine, adenosine receptors and their role in glucose homeostasis and lipid metabolism. J Cell Physiol 2013.

26. Allard B, Allard D, Buisseret L, Stagg J. The adenosine pathway in immuno-oncology. Nat Rev Clin Oncol 2020.

27. Willingham SB, Criner G, Hill C, Hu S, Rudnick JA, Daine-Matsuoka B, Hsieh J, et al. Characterization and Phase 1 Trial of a B Cell Activating Anti-CD73 Antibody for the Immunotherapy of COVID-19. medRxiv 2020:2020.2009.2010.20191486.

28. Tabula Muris C, Overall c, Logistical c, Organ c, processing, Library p, sequencing, et al. Single-cell transcriptomics of 20 mouse organs creates a Tabula Muris. Nature 2018;562:367–372.

29. Ang CH, Hsu SH, Guo F, Tan CT, Yu VC, Visvader JE, Chow PKH, et al. Lgr5(+) pericentral hepatocytes are self-maintained in normal liver regeneration and susceptible to hepatocarcinogenesis. Proc Natl Acad Sci U S A 2019;116:19530–19540.

30. Wang B, Zhao L, Fish M, Logan CY, Nusse R. Self-renewing diploid Axin2(+) cells fuel homeostatic renewal of the liver. Nature 2015;524:180–185.

31. Gwinn DM, Shackelford DB, Egan DF, Mihaylova MM, Mery A, Vasquez DS, Turk BE, et al. AMPK phosphorylation of raptor mediates a metabolic checkpoint. Mol Cell 2008;30:214–226.

32. Pastor-Anglada M, Perez-Torras S. Who Is Who in Adenosine Transport. Front Pharmacol 2018;9:627.

33. Cai Y, Li H, Liu M, Pei Y, Zheng J, Zhou J, Luo X, et al. Disruption of adenosine 2A receptor exacerbates NAFLD through increasing inflammatory responses and SREBP1c activity. Hepatology 2018;68:48–61.

34. Garcia D, Hellberg K, Chaix A, Wallace M, Herzig S, Badur MG, Lin T, et al. Genetic Liver-Specific AMPK Activation Protects against Diet-Induced Obesity and NAFLD. Cell Rep 2019;26:192–208 e196.

35. Volmer JB, Thompson LF, Blackburn MR. Ecto-5’-nucleotidase (CD73)-mediated adenosine production is tissue protective in a model of bleomycin-induced lung injury. J Immunol 2006;176:4449–4458.

36. Grenz A, Zhang H, Eckle T, Mittelbronn M, Wehrmann M, Kohle C, Kloor D, et al. Protective role of ecto-5’-nucleotidase (CD73) in renal ischemia. J Am Soc Nephrol 2007;18:833–845.

37. Gehrke N, Schattenberg JM. Metabolic Inflammation-A Role for Hepatic Inflammatory Pathways as Drivers of Comorbidities in Nonalcoholic Fatty Liver Disease? Gastroenterology 2020;158:1929–1947 e1926.

38. Mauvais-Jarvis F, Bairey Merz N, Barnes PJ, Brinton RD, Carrero JJ, DeMeo DL, De Vries GJ, et al. Sex and gender: modifiers of health, disease, and medicine. Lancet 2020;396:565–582.

39. Heron M. Deaths: Leading Causes for 2017. Natl Vital Stat Rep 2019;68:1–77.

40. Vilar-Gomez E, Calzadilla-Bertot L, Wai-Sun Wong V, Castellanos M, Aller-de la Fuente R, Metwally M, Eslam M, et al. Fibrosis Severity as a Determinant of Cause-Specific Mortality in Patients With Advanced Nonalcoholic Fatty Liver Disease: A Multi-National Cohort Study. Gastroenterology 2018;155:443–457 e417.

41. Lonardo A, Nascimbeni F, Ballestri S, Fairweather D, Win S, Than TA, Abdelmalek MF, et al. Sex Differences in Nonalcoholic Fatty Liver Disease: State of the Art and Identification of Research Gaps. Hepatology 2019;70:1457–1469.

42. Peralta C, Bartrons R, Serafin A, Blazquez C, Guzman M, Prats N, Xaus C, et al. Adenosine monophosphate-activated protein kinase mediates the protective effects of ischemic preconditioning on hepatic ischemia-reperfusion injury in the rat. Hepatology 2001;34:1164–1173.

43. Mulligan JD, Gonzalez AA, Kumar R, Davis AJ, Saupe KW. Aging elevates basal adenosine monophosphate-activated protein kinase (AMPK) activity and eliminates hypoxic activation of AMPK in mouse liver. J Gerontol A Biol Sci Med Sci 2005;60:21–27.

44. Hwang AB, Ryu EA, Artan M, Chang HW, Kabir MH, Nam HJ, Lee D, et al. Feedback regulation via AMPK and HIF-1 mediates ROS-dependent longevity in Caenorhabditis elegans. Proc Natl Acad Sci U S A 2014;111:E4458–4467.

45. Rabinovitch RC, Samborska B, Faubert B, Ma EH, Gravel SP, Andrzejewski S, Raissi TC, et al. AMPK Maintains Cellular Metabolic Homeostasis through Regulation of Mitochondrial Reactive Oxygen Species. Cell Rep 2017;21:1–9.

46. Burnstock G, Vaughn B, Robson SC. Purinergic signalling in the liver in health and disease. Purinergic Signal 2014;10:51–70.

47. Hotamisligil GS. Inflammation, metaflammation and immunometabolic disorders. Nature 2017;542:177–185.

48. Ohta A, Sitkovsky M. Role of G-protein-coupled adenosine receptors in downregulation of inflammation and protection from tissue damage. Nature 2001;414:916–920.

49. Chouker A, Thiel M, Lukashev D, Ward JM, Kaufmann I, Apasov S, Sitkovsky MV, et al. Critical role of hypoxia and A2A adenosine receptors in liver tissue-protecting physiological anti-inflammatory pathway. Mol Med 2008;14:116–123.

50. Cordero-Herrera I, Kozyra M, Zhuge Z, McCann Haworth S, Moretti C, Peleli M, Caldeira-Dias M, et al. AMP-activated protein kinase activation and NADPH oxidase inhibition by inorganic nitrate and nitrite prevent liver steatosis. Proc Natl Acad Sci U S A 2019;116:217–226.

## References

1. Postic C, Shiota M, Niswender KD, Jetton TL, Chen Y, Moates JM, Shelton KD, et al. Dual roles for glucokinase in glucose homeostasis as determined by liver and pancreatic beta cell-specific gene knock-outs using Cre recombinase. J Biol Chem 1999;274:305–315.

2. Fu D, Lippincott-Schwartz J. Monitoring the Effects of Pharmacological Reagents on Mitochondrial Morphology. Curr Protoc Cell Biol 2018;79:e45.

3. Snider NT, Griggs NW, Singla A, Moons DS, Weerasinghe SV, Lok AS, Ruan C, et al. CD73 (ecto-5’-nucleotidase) hepatocyte levels differ across mouse strains and contribute to mallory-denk body formation. Hepatology 2013;58:1790–1800.

